# Gut microbiome structure and adrenocortical activity in dogs with aggressive and phobic behavioral disorders

**DOI:** 10.1101/573865

**Authors:** E Mondo, M Barone, M Soverini, F D’Amico, M Cocchi, C Petrulli, M Mattioli, G Marliani, M Candela, PE Accorsi

## Abstract

Accompanying human beings since the Paleolithic period, dogs has been recently regarded as a reliable model for the study of the gut microbiome connections with health and disease. In order to provide some glimpses on the connections between the gut microbiome layout and host behavior, we profiled the phylogenetic composition and structure of the canine gut microbiome of dogs with aggressive (n = 17), phobic (n = 15) and normal behavior (n = 17). According to our findings, aggressive behavioral disorder was found to be characterized by a peculiar gut microbiome structure, with high biodiversity and enrichment in generally subdominant bacterial genera. On the other hand, phobic dogs were enriched in *Lactobacillus*, a bacterial genus with known probiotic and psychobiotic properties. Although further studies are needed to validate our findings, our work supports the intriguing opportunity that different behavioral phenotypes in dogs may be associated with peculiar gut microbiome layouts, suggesting possible connections between the gut microbiome and the central nervous system and indicating the possible adoption of probiotic interventions aimed at restoring a balanced host-symbiont interplay for mitigating behavioral disorders.

## Introduction

Descending from the gray wolf (*Canis lupus*), dogs were domesticated during the Paleolithic period, accompanying humans across the transition from hunting-gathering to rural agriculture of the Neolithic, to post-industrialized Western lifestyle [1–3]. The frequent sharing of food resources with human beings has been a selective force able to drive changes in the digestive and metabolic system of dogs, enabling them to efficiently adapt to a more starch-enriched diet compared to their wild ancestor, and ultimately influence canine behavior [4,5]. The canine gastrointestinal tract harbors a complex and highly biodiverse microbial ecosystem, whose predominant taxa resemble those typically found in the gut of other omnivorous mammals. However, in comparison to both mice and pigs, the canine gut microbiome (GM) result the most similar to humans [6,7]. Thus, in dogs, the GM-host mutualistic exchange well approximate what has been observed in humans [8–11]. Indeed, the peculiar patterns of dysbiosis observed in dogs with IBD are generally comparable to variations typically found in humans, suggesting that bacterial responses to inflammatory conditions are conserved among the two [12,13]. As observed in humans, the eubiotic and stable configuration of the canine GM is therefore of fundamental importance for the maintenance of a homeostatic gut environment and of the overall host health.

Several recent studies have shown the ability of the mammalian GM to communicate with the host central nervous system (CNS) through several parallel channels, involving the vagus nerve, neuroimmune and neuroendocrine signaling mechanisms, and the production of neuroactive chemicals – i.e. gamma-aminobutyric acid (GABA), serotonin (5-HT), norepinephrine and dopamine [14–17]. Conversely, the CNS can influence GM structure and metabolome, influencing the gut environment, acting on motility, secretion and permeability via the autonomic nervous system (ANS) [18]. It is thus a matter of fact that the GM can influence the host behavior and *vice versa*, exerting a key role in the modulation of the gut-brain axis. The development of researches during the last decades indeed suggests the presence of a bidirectional communication between gut and brain.

Despite the high variability and severity of behavioral disorders observed in dogs, the aggressive behavior has been found to be the most common, followed by separation anxiety and phobia [19]. Aggressive, anxiety and phobia behavioral disorders can be considered as stress responses with an increase in glucocorticoid (GC) secretion mediated by the hypothalamic-pituitary-adrenal (HPA) axis [20,21]. Although recent works on canine microbiome have investigated potential interactions with aggression, these studies have focused on the variations of their GM profile after targeted dietary interventions to reduce aggressive behaviors [22,23]. To the best of our knowledge, no study has focused on the comparison of the GM structure between dogs exhibiting aggressive, phobic and normal behavior, with specific associations with adrenocortical activity. In order to provide some glimpses in this direction, in our work, 49 dogs – 31 males and 18 females – of different breed, age, and weight, housed in individual boxes were enrolled. Following the behavioral evaluation performed by a behaviorist veterinary and a dog handler, dogs were classified into three groups: aggressive, phobic and normal behavior. We then profiled the phylogenetic composition and structure of their canine GM, measuring the levels of fecal hormones (i.e. cortisol and testosterone) to evaluate the adrenocortical activity in each study group. Fecal GC and their metabolites may constitute a reliable non-invasive proxy of adrenocortical activity, reflecting long-term stress responses with less interference from acute stressors [24–26].

Our data suggest that peculiar microbiome configurations could be connected with dogs’ behavioral phenotype, which in turn might be exacerbated by altered proportions of gut symbionts, ultimately triggering a feedback loop. In this scenario, targeted interventions on GM could constitute a valuable tool to restore a proper host-symbiont interplay, finally mitigating behavioral disorders and improving the overall host health.

## Results

### Relative abundance of major gut microbiome components in the enrolled cohort

In order to evaluate possible differences in GM communities among the behavioral study groups, we collected fecal samples from 49 dogs. In particular, 31 males and 18 females of different breed, aged between 1 and 13 years, were recruited from three animal shelters located in the metropolitan area of Bologna (Italy). Following the behavioral evaluation performed by a behaviorist veterinary and a dog handler, dogs were grouped based on their behavioral phenotype: 17 were classified as aggressive and 15 as phobic, while 17 exhibited a normal behavior. The 16S rRNA sequencing, performed using the Illumina MiSeq platform, yielded a total of 1,611,153 high-quality sequences, with an average of 32,880 ± 6,009 sequences per sample (range 20,664 – 45,486), subsequently clustered in 8,460 OTUs with a 97% identity threshold.

The most abundant phyla detected within the normal behavior samples were Firmicutes (68.0 ± 4.6%), Bacteroidetes (13.7 ± 3.6%), and Actinobacteria (9.9 ± 1.6%), with Fusobacteria (4.8 ± 1.3%) and Proteobacteria (2.1 ± 0.8%) as minor components. The aggressive group showed similar proportions among the dominant phyla when compared to the normal behavior group, except for a reduced relative abundance of Bacteroidetes (P-value = 0.02; Wilcoxon test). No significant differences at phylum level were detected between the phobic and the normal behavior groups, as well as between the phobic and the aggressive groups.

At family level, Lachnospiraceae, Erysipelotrichaceae and Clostridiaceae constitute the major components of the normal behavior group (relative abundance > 10%). A depletion in the relative abundance of Bacteriodaceae, Alcaligenaceae and [Paraprevotellaceae], as well as an increase in Erysipelotrichaceae (P-value < 0.05) was observed in the aggressive group compared to the normal behavior group. The phobic group was instead characterized by an increase in the relative abundance of the family Rikenellaceae (P-value = 0.04) when compared to the normal behavior group. Aggressive and phobic groups were found to be distinguishable due to different proportions in the relative abundance of the bacterial families [Mogibacteriaceae] and Veillonellaceae, respectively depleted and enriched in the aggressive group (P < 0.04) (Fig 1A, 1B).

**Fig 1.**
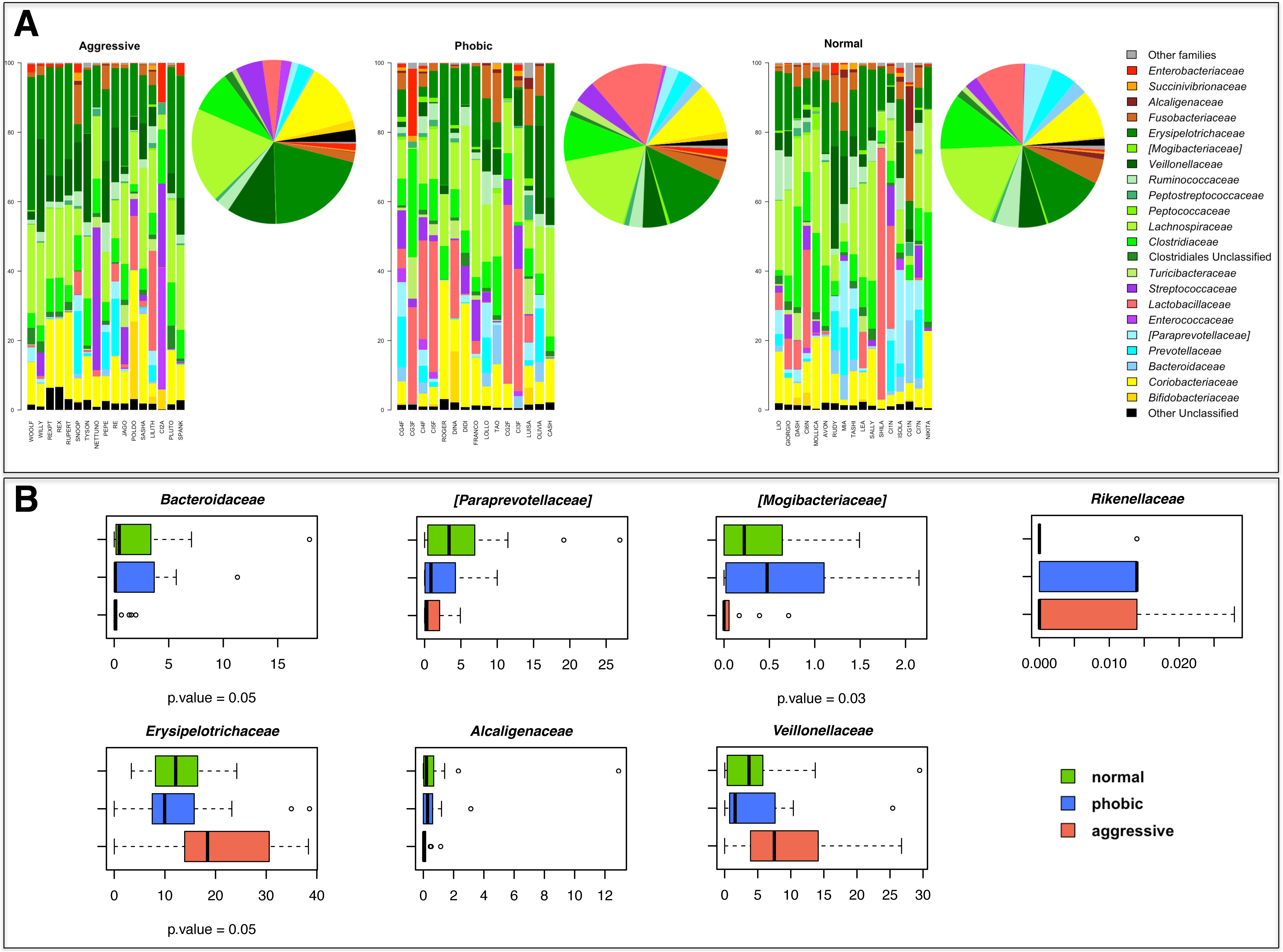
Canine gut microbiome profile of the behavior groups. (A) Relative abundances of family-level taxa in each subject of the enrolled cohort (barplots) and respective average values of each study group (piecharts). (B) Boxplots showing the distribution of the relative abundances of bacterial families enriched or depleted within the gut microbiota of aggressive or phobic groups.

At the genus taxonomic level, *Clostridium*, *Lactobacillus*, *Blautia* and *Collinsella* represent the major portion of the normal behavior group GM (relative abundance > 5%). Several microbial genera were found to be significantly depleted in the aggressive group. The relative abundance of the genera *Oscillospira*, *Peptostreptococcus*, *Bacteroides*, *Sutterella*, and *Coprobacillus* were significantly lower in aggressive compared to normal behavior group, while *Catenibacterium*, *Megamonas* and [*Eubacterium*] showed an opposite trend (P-value < 0.04). At the genus level, no significant differences were detected between the phobic and the normal behavior group. The differences found between phobic and aggressive groups were due to an increased relative abundance of *Catenibacterium* and *Megamonas* in the latter group (P < 0.007), in addition to a slightly depletion of the genus *Epulopiscium* (P = 0.04) (S3 Fig).

### Comparison of the overall gut microbiome compositional structure between aggressive, phobic and normal behavior dogs

The intra-individual diversity of the canine GM was assessed by means of the phylogenetic metric Observed OTUs and the Shannon biodiversity index at the genus level. While according to the Shannon index the intra-individual GM diversity was comparable between aggressive, phobic and normal behavior groups (mean ± SD, 5.2 ± 0.8, 5.14 ± 0.7, and 5.2 ± 0.7, respectively), the Observed OTUs metric highlighted a statistically significant difference between aggressive and phobic groups (P-value = 0.02; Kruskal-Wallis test). In particular, the aggressive group was characterized by the higher number of different taxa observed (mean ± SD, 583.9 ± 310.6), the phobic group had the lower value (430.07 ± 112.28), and the normal behavior group exhibited intermediate values (454.8 ± 118.3) (Fig 2).

**Fig 2.**
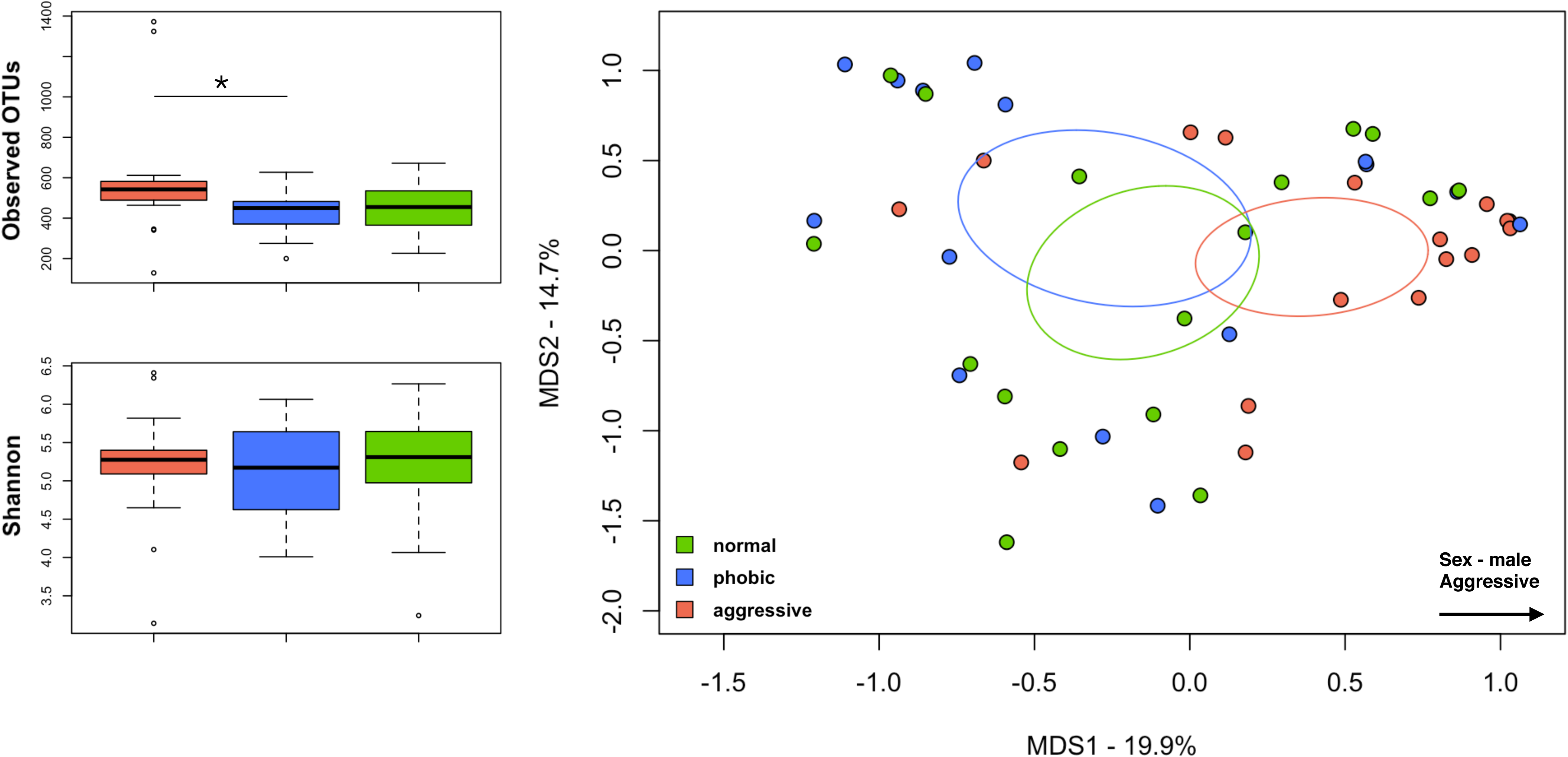
Biodiversity of the canine gut microbiome. Boxplot showing the alpha diversity measures computed with phylogenetic and non-phylogenetic metrics (Shannon diversity index, observed OTUs). Behavior-related groups are identified with colored box and whiskers (orange, Aggressive; blue, Phobic; green, Normal). Significant difference was found between aggressive and phobic groups, according to the observed OTUs metric (P-value = 0.02; Kruskal-Wallis test). Principal coordinate analysis (PCoA) plots showing the beta diversity of the intestinal bacterial communities of the study groups, based on Jaccard similarity index. A significant separation between aggressive and phobic behavior groups was found (P-value = 0.02, permutation test with pseudo-*F* ratios).

According to the Jaccard similarity index, the Principal Coordinates Analysis (PCoA) of the inter-sample variation highlighted a significant separation between the structural composition of the GM among study groups (P-value = 0.02, permutation test with pseudo-*F* ratios) (Fig 2). In order to identify bacterial drivers that contribute to groups clustering (permutation correlation test, P-value < 0.001), a superimposition of the genus relative abundance was performed on the PCoA plot. As showed in S1 Fig, major drivers of the normal behavior group segregation were *Faecalibacterium*, *Bacteroides*, *Phascolarctobacterium, Fusobacterium*, *Prevotella*, and [*Prevotella*]. The phobic group was characterized by an enrichment in *Lactobacillus*, while *Dorea*, *Blautia*, *Collinsella*, [*Ruminococcus*], *Slackia*, *Catenibacterium*, and *Megamonas* were more represented within the aggressive group.

To dissect discriminant GM component for dogs showing aggressive behavior, we applied the machine learning method Random Forest [27] to the genus level data set. Behavior-discriminatory bacterial genera were identified with distinctive changes in their relative abundances (Fig 3). Specifically, our analysis revealed two ‘aggressive-discriminatory’ bacterial genera: *Catenibacterium* and *Megamonas*.

**Fig 3.**
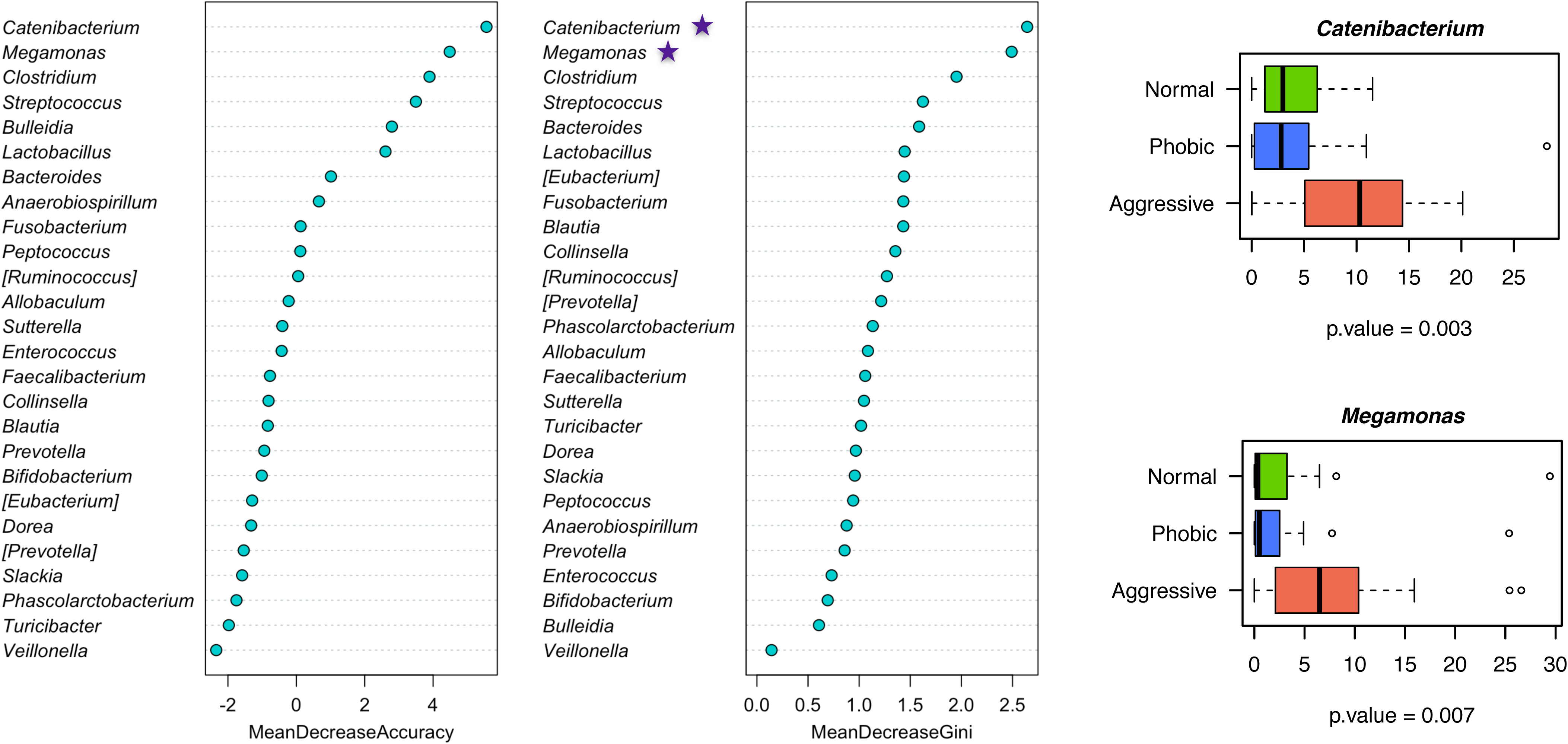
Behavior-related gut microbiome signature. Top 26 features from the obtained dataset as revealed by Random Forest. Stars denote the bacterial genera discriminant of aggressive group. Boxplots shows the comparison of the relative abundances of these bacterial genera between the study groups.

### Fecal cortisol and testosterone levels

Cortisol and testosterone levels were measured in fecal samples through RIAs. The statistical analysis did not evidence any significant differences between the three groups of dogs for hormonal data. Our results showed that the three study groups had similar median value of fecal cortisol and testosterone levels (Fig 4A). However, it should be noticed that the range of testosterone level in aggressive and phobic populations is largest than in normal, so we can assume that there is a greater variability between subjects (Fig 4B). As consequences of the previously described results, the median of testosterone/cortisol (T/C) ratio were similar in all the three groups, only slightly higher in phobic than in aggressive and normal dogs (Fig 4C).

**Fig 4.**
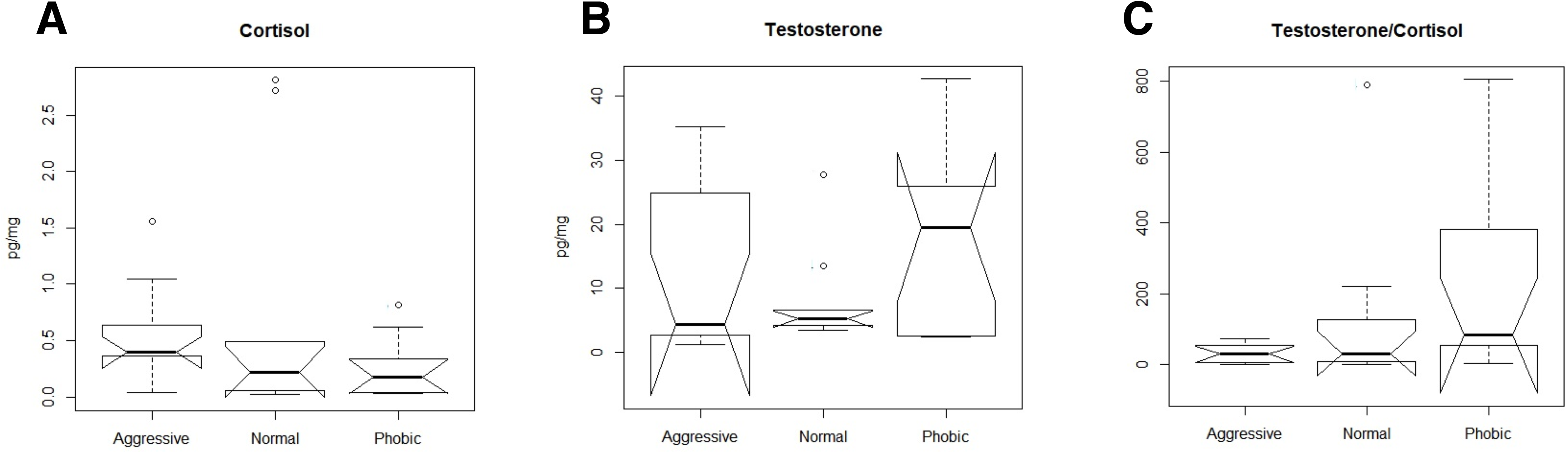
Fecal hormone levels of the behavior groups. Boxplots showing levels of cortisol (A) and testosterone (B), and testosterone/cortisol ratio (C) detected in stool samples of the study groups. No significant difference was found among study groups (P-value > 0.05, Kruskal-Wallis test).

## Discussion

Within the present work, we profiled the GM structure and measured the fecal cortisol and testosterone levels of 49 dogs - 31 males and 18 females - of different breed, age, and weight, housed in individual boxes of three animal shelters located in the metropolitan area of Bologna (Italy). Dogs were classified into three study groups based on their behavioral phenotype: aggressive, phobic or normal behavior. The phylogenetic profiles of the canine GM observed in our cohort were found to be in line with those already reported in literature for healthy dogs [9,28,29], but with a slightly higher abundance of Firmicutes and Actinobacteria, as well as a corresponding lower abundance of Bacteroidetes and Proteobacteria. According to our results, the aggressive group GM is characterized by a higher number of observed OTUs compared to both phobic and normal behavior groups. Interestingly, the GM structure of our cohort segregate according to the behavioral disorder of the host, showing a stronger separation of the aggressive group. The latter group seems to be defined by a higher abundance of typically subdominant taxa, such as *Dorea*, *Blautia*, *Collinsella*, [*Ruminococcus*], *Slackia*, *Catenibacterium*, and *Megamonas*. Conversely, the phobic group is characterized by an enrichment of *Lactobacillus*, a bacterial genus comprising well-known GABA producers, the main CNS inhibitory neurotransmitter able to regulate emotional behavior in mice via the vagus nerve [30,31]. The major drivers for the normal behavior group segregation are *Faecalibacterium*, *Bacteroides*, *Phascolarctobacterium*, *Fusobacterium*, *Prevotella* and [*Prevotella*], reflecting the predominance of bacterial genera commonly associated with the GM of healthy dogs [12,32]. Finally, according to the literature, no correlation was observed regarding the phobic behavior in relation to sex or age [19].

According to our results, fecal cortisol and testosterone levels of aggressive dogs did not significantly differ from those of phobic and normal dogs. Aggressive dogs are well known to possess higher blood concentrations of cortisol and lower serotonin levels than non-aggressive dogs [20]. However, fecal cortisol levels are not influenced by the activation of HPA axis during the sampling procedure, which is itself stressful for the animal [33]. This can possibly explain the differences in the observed cortisol levels between studies carried out in blood or fecal samples, Testosterone is often correlate with aggressive behaviors in many species [34], but this association is not completely demonstrated in dogs. Indeed, some studies have evidenced that the castration reduce only mildly aggressiveness and that neutered dogs can be more aggressive [35,36]. In contrast with results of Terburg *et al.* [37], who suggested that a high value of T/C ratio may be a predictive factor of aggressive behavior, we found no significant differences between groups.

Studies about cortisol in phobic state are confusing, and it seems that cortisol does not increase, or increase only slightly in phobic subjects compared to normal ones [38]. Our results did not show any significant differences between phobic and normal dogs for cortisol concentration. However, the cortisol level of phobic dogs is slightly lower than normal ones. Phobic state can cause a chronic and excessive stress [39], and scientific literature about phobic dogs suggests that they can tend to a depressive state. In human beings and in dogs, depression is characterized by lower cortisol and serotonin levels [40].

Our results suggest that dysbiotic GM configurations in a long-term stress levels scenario might influence the local gut environment through the release of potentially neuroactive microbial by-products, probably affecting the behavior of the host mainly as a side effect. In particular, dogs exhibiting aggressive behavioral disorders were characterized by a peculiar GM structure, a high biodiversity and an enrichment in generally subdominant bacterial genera. We then applied a machine learning method (Random Forest) to our genus level data set, identifying *Catenibacterium* and *Megamonas* as bacterial discriminants of aggressive behavior. Respectively belonging to the families Erysipelotrichaceae and Veillonellaceae, *Catenibacterium* and *Megamonas* have been recently correlated with primary bile acid metabolism and abdominal pain in humans [41–43], suggesting a possible connection between a dysbiotic GM profile and behavioral disorders. We found no alteration within the GM of dogs with phobic behavioral disorder except for an increase in *Lactobacillus*, bacterial genera with well-known probiotic properties. Interestingly, chronic treatment with *L.rhamnosus* can influence anxiety- and depression-related behavior by modulating GABA receptor mRNA expression in specific brain regions [30]. Even if it is impossible to dissect the factors supporting the increase of *Lactobacillus* observed in phobic dogs, it is tempting to speculate that a higher abundance of this psychobiotics [44,45] could contribute to the establishment of peculiarities of the phobic behavioral phenotype. However, further studies are required to validate our findings. Indeed, the limitations due to the small number of enrolled animals imply a limited statistical power. Further, invasive blood sampling will be necessary, allowing to match blood and fecal cortisol and testosterone levels to find robust connections with the GM state. Nonetheless, our study support the intriguing opportunity that different behavioral phenotypes in dogs associate with peculiar GM layouts. Particularly, aggressive dogs possess dysbiotic GM configuration, possibly exacerbating the host aggressiveness and supporting the adoption probiotic interventions aimed at restoring a balanced host-symbiont interplay, improving the overall host health and eventually mitigating behavioral disorders. This preliminary research can thus be considered a starting point for future studies of clinical interest, deepening the understanding of the mechanisms underlying the relationship between behavioral disorders and the GM, ultimately providing new insight into veterinary behavioral medicine and facilitating a predictive diagnosis of canine behavior.

## Materials and methods

### Enrolled animals, behavioral evaluation and sample collection

The entire study was previously evaluated and approved by the Scientific Ethic Committee for Animal Experimentation (University of Bologna). All the procedures were monitored by the responsible of the Department of Veterinary Medical Science (DIMEVET) for animal welfare.

In the study, 49 dogs (31 males and 18 females) of different breed, age, and weight, housed in individual boxes of three different animal shelters located in the metropolitan area of Bologna (Italy) were enrolled (S1 Table). The animals were fed on a mixed diet, including wet and dry commercial feed and additional homemade food.

For each animal, a behavioral evaluation was performed by a behaviorist veterinary and a dog handler, classifying the dogs enrolled based on their behavioral phenotype. Seventeen animals were classified as aggressive and 15 as phobic, while 17 animals exhibited a normal behavior.

Fecal samples were collected from each animal immediately after the evacuation, between 7:00 and 12:00 am, avoiding debris and cross-contaminations. Specimens were frozen with liquid nitrogen and transported to the laboratory, then stored at −80°C until DNA extraction and sequencing. The remaining part of fecal samples was picked up through non-sterile bags and frozen at −20°C until cortisol and testosterone assay.

### Fecal cortisol and testosterone radioimmunoassays

Cortisol and testosterone concentrations were determined by radioimmunoassays (RIAs) based on binding of ^3^H-steroid by competitive adsorption [46]. All concentrations were expressed in pg/mg of fecal matter. Extraction methodology was modified from Schatz & Palme [21]. Cortisol and testosterone were extracted from fecal specimens (500 mg, wet weight) with methanol-water solution (5 ml, v/v 4:1) and ethyl ether (5 ml). The portion of ether was vaporized under an airstream suction hood at 37° C. Dry residue was finally dissolved again into 0.5 ml PBS (0.05 M, pH 7.5), adding 125, 250, 500, or 1000 pg of ^3^H-cortisol or ^3^H-testosterone to 500 mg of feces and incubating for 30 min at room temperature, performing a recovery test on five replicates. The extraction was performed as described before, yielding a mean percentage recovery of 87.5 ± 2.4 and 89.3 ± 2.1 for cortisol and testosterone, respectively. Cortisol and testosterone metabolites assay in feces were carried out according to Tamanini [47] and Gaiani [48], respectively. Analysis were performed in duplicates. The cortisol RIA was performed using an antiserum to cortisol-21-hemisuccinate-BSA (anti- rabbit), at a working dilution of 1:20 000 and ^3^H-cortisol (30 pg/tube vial) as tracer. The testosterone RIA was performed using an antiserum to testosterone-3-carboxymethyloxime-BSA (anti- rabbit), at a working dilution of 1:35 000 and ^3^H-testosterone (31 pg/tube vial) as tracer. Fecal cortisol and testosterone samples containing high concentrations of endogenous steroids (100 µl) were serially diluted through PBS (0.05 M, pH 7.5) in volumes of 50, 25, 10 and 5 ml, in order to determine the parallelism between cortisol and testosterone standards. Parallelism was assessed between these serial dilutions of standards (ranging from 7.8 to 1000 pg/100 ml tube vial). Validation parameters of analysis were: sensitivity 0.19 pg/mg, intra-assay variability 5.9%, inter-assay variability 8.7%, for cortisol; sensitivity 1.1 pg/mg, intra-assay variability 6.2%, inter-assay variability 9.6%, for testosterone. Radioactivity was determined using a liquid scintillation β counter and a linear standard curve, ad hoc designed by a software program [49].

### Bacterial DNA extraction from stool samples

Total microbial DNA was extracted from each fecal sample using the DNeasy Blood & Tissue kit (QIAGEN), with the modified protocol described by Turroni *et al.* [50]. Briefly, 250 mg of feces were resuspended in 1 ml of lysis buffer (500 mM NaCl, 50 mM Tris-HCl pH 8, 50 mM EDTA, 4% SDS). Fecal samples were added with four 3-mm glass beads and 0.5 g of 0.1-mm zirconia beads (BioSpec Products, Bartlesville, USA) and homogenized with 3 bead-beating steps using the FastPrep instrument (MP Biomedicals, Irvine, CA) at 5.5 movements/s for 1 min, keeping the samples on ice for 5 min after each treatment. Samples were subsequently heated at 95°C for 15 min and centrifuged to pellet stool particles. Supernatants were added with 260 μl of 10 M ammonium acetate, centrifuged for 10 min at full speed, and incubated in ice for 30 min with one volume of isopropanol. Nucleic acids were collected by centrifugation, washed with 70% ethanol and resuspended in 100 μl of TE buffer. RNA and protein removal was performed by incubating the samples with DNase-free RNase (10 mg/ml) at 37°C for 15 min and protease K at 70°C for 10 min, respectively. Subsequently, DNA purification with QIAmp Mini Spin columns were performed as per manufacturers instruction. The extracted bacterial DNA was quantified using the NanoDrop ND-1000 spectrophotometer (NanoDrop Technologies).

### PCR amplification and sequencing

The V3-V4 region of the 16S rRNA was amplified with PCR using 200 nmol/l of S-D-Bact-0341-b-S-17/S-D-Bact-0785-a-A-21 primers [51] with Illumina overhang adapter sequences, in a final volume of 25 μl containing 12.5 ng of genomic DNA and 2X KAPA HiFi HotStart ReadyMix (Kapa Biosystems). PCR reactions were performed in a Thermal Cycle T gradient (Biometra GmbH) using the following thermal program: 3 min at 95°C for the initial denaturation, followed by 25 cycles of denaturation at 95°C for 30 sec, annealing at 55°C for 30 sec, extension at 72°C for 30 sec, and a final extension step at 72°C for 5 min. PCR products of about 460 bp were purified using a magnetic bead-based system (Agencourt AMPure XP; Beckman Coulter) and sequenced on Illumina MiSeq platform using the 2 x 250 bp protocol, according to the manufacturer’s instructions (Illumina). The libraries were pooled at equimolar concentrations, denatured and diluted to 6 pmol/l before loading onto the MiSeq flow cell.

### Bioinformatics and statistics

Raw sequences were processed using a pipeline combining PANDAseq [52] and QIIME [53]. The UCLUST software [54] was used to bin high-quality reads into operational taxonomic units (OTUs) through an open-reference strategy at a 0.97 similarity threshold. Taxonomy was assigned using the RDP classifier and the Greengenes database as a reference (release May 2013). Chimera filtering was performed discarding all singleton OTUs. Alpha rarefaction was evaluated by using the Observed OTUs metric, and the Shannon biodiversity index, which aims to measure diversity based on evenness, while beta diversity was estimated according to the Jaccard similarity. Random Forests and all statistical analysis was computed using R software (version 3.1.3) and the packages randomForest, vegan and made4. The significance of data separation on the PCoA was tested using a permutation test with pseudo-*F* ratios (function adonis of vegan package). Non-parametric and correlation tests were achieved with Wilcoxon rank-sum or Kruskal-Wallis test and the Kendall tau, respectively. Cortisol and testosterone concentrations, as well as T/C ratio was analyzed using the normality test of Shapiro-Wilk, in order to establish the distribution of each variable in the population. P-values < 0.05 were considered statistically significant.

## Supporting information

**S1 Table. Metadata of the enrolled cohort.**

**S1 Fig. Superimposition of the genus relative abundance on the PCoA plot.** Arrows represent the direction of significant correlations (permutation correlation test, P-value < 0.001).

**S2 Fig. Main bacterial genera represented within the canine gut microbiome.**

