## Supplementary figures and images for "Gut microbiome structure and adrenocortical activity in dogs with aggressive and phobic behavioral disorders"

### Supplemental Figure 1

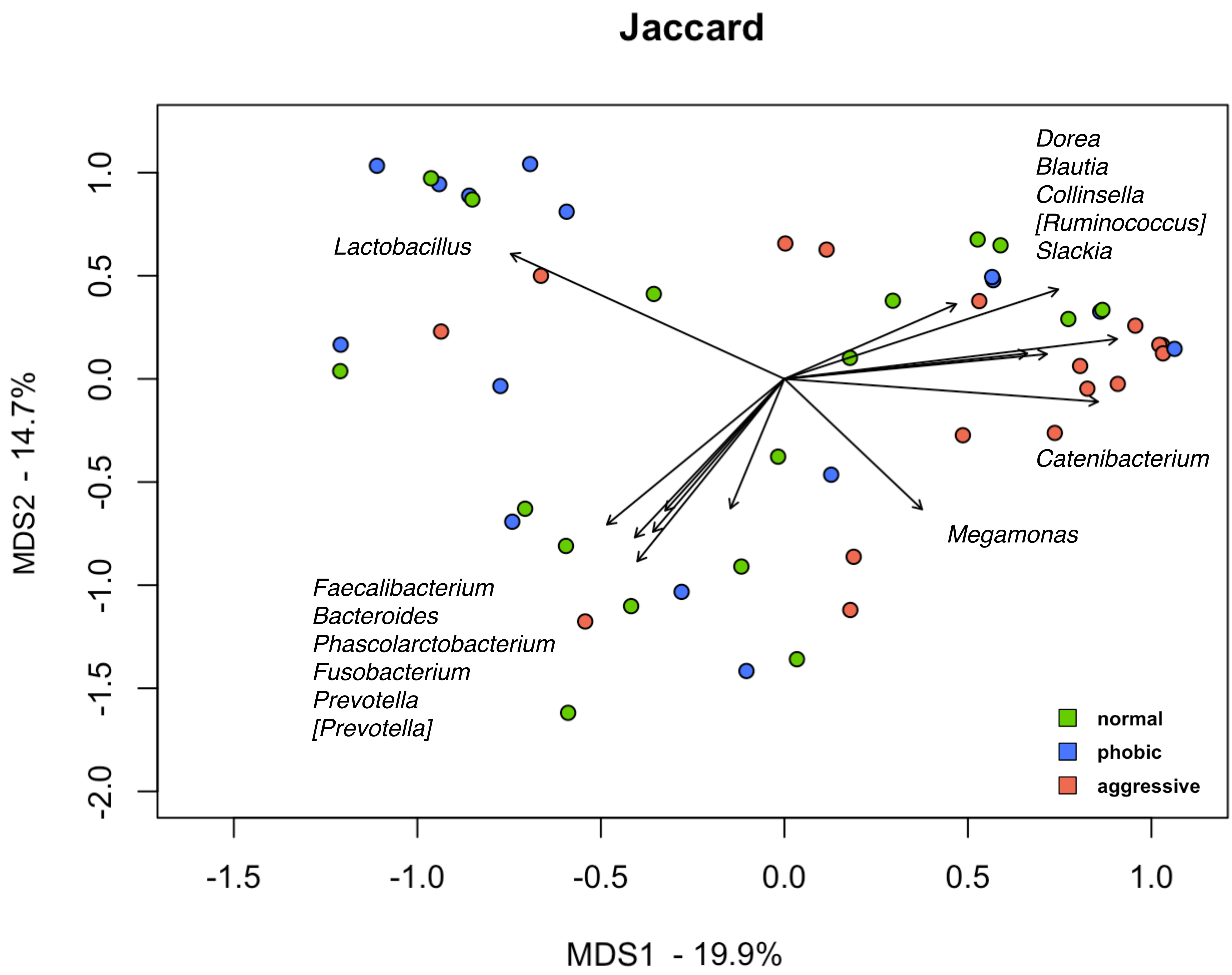

### Supplemental Figure 2

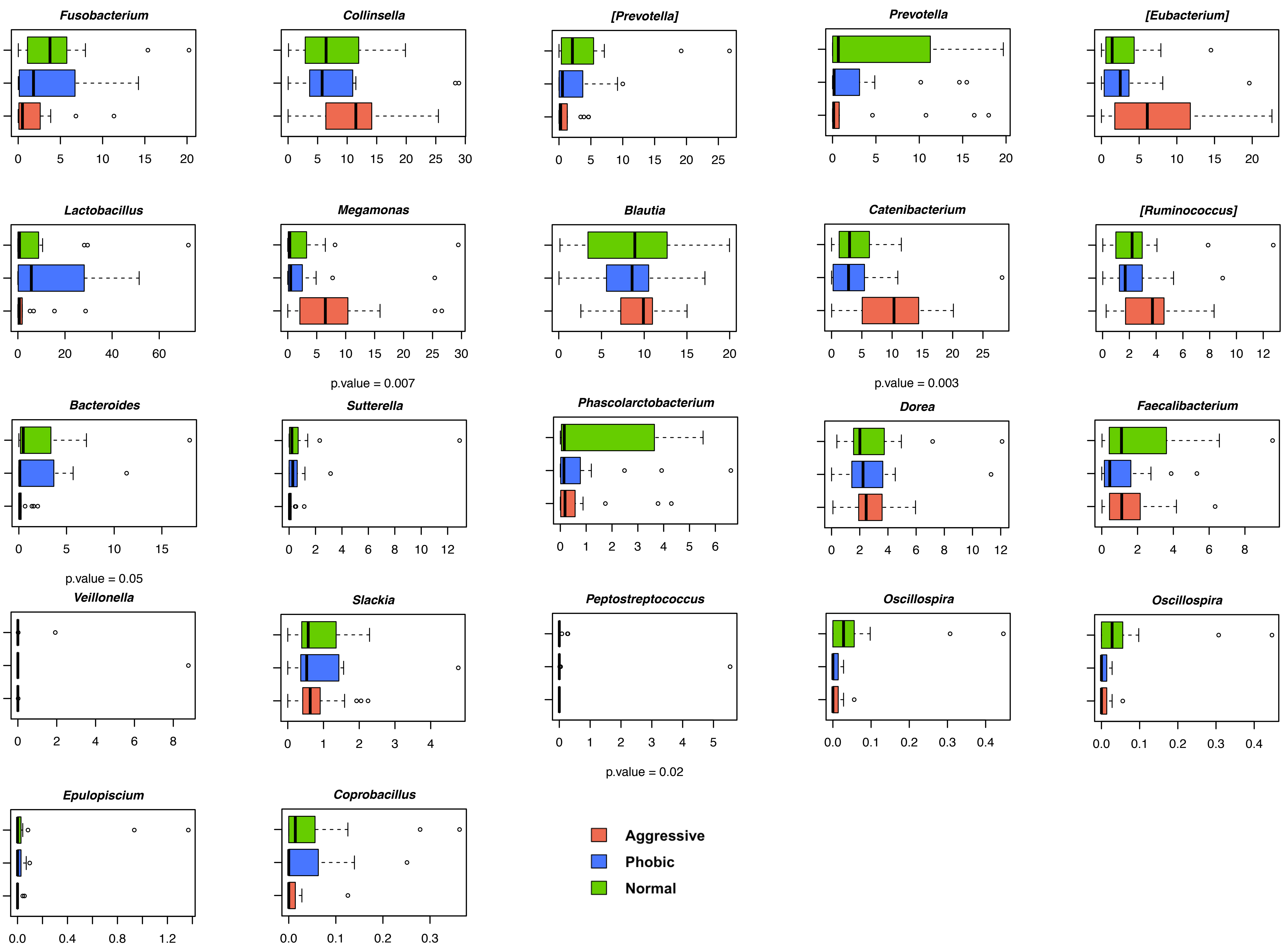
